# A Hotspot Atop: Rivers of the Guyana Highlands Hold High Diversity of Endemic Pencil Catfish

**DOI:** 10.1101/640821

**Authors:** Holden J. Paz, Malorie M. Hayes, Carla C. Stout, David C. Werneke, Jonathan W. Armbruster

## Abstract

**Aim:** The Pakaraima Mountains are an ancient mountain range along the borders of Guyana, Brazil, and Venezuela. The high plateau is drained by multiple river systems in all directions. Although hypotheses have been presented for the biogeographic relationships of lowland rivers, the interconnectivity of rivers on the top of the plateau is unknown. With multiple complex rivers in a small, upland area we predicted a high level of endemism for stream fishes and complex biogeographic relationships. We explore this with the incredibly diverse pencil catfish genus *Trichomycterus*. Only two species are known from the region. In this study, we 1) confirm the discovery of multiple endemic *Trichomycterus* species in the region, 2) determine the phylogenetic placement of our samples to posit biogeographical scenarios, and 3) provide clarification for the identification of *T. guianensis* based on morphology.

**Location:** Pakaraima Mountains, a part of the Guiana Shield in Guyana, South America

**Taxon:** Pencil catfish genus *Trichomycterus*

**Methods:** Using collections from recent expeditions to the Pakaraima Mountains of Guyana, we amplified three mitochondrial (16S, COI, and cytb) and two nuclear markers (myh6 and RAG2). We constructed individual gene trees as well as a concatenated tree to determine the placement of these taxa within the *Trichomycterus* of the Trans-andean/Amazonian clade.

**Results:** Our results identify six unique lineages in the highlands of Guyana. Only two species, *Trichomycterus guianensis* and *T. conradi*, were previously known to science.

**Main Conclusions:** The Pakaraima Mountains of South America are a region of high endemism, as demonstrated here in *Trichomycterus* catfishes. We find two species occupying multiple basins, suggesting that Pakaraima streams either maintain or had some degree of recent connectivity. We identify six endemic lineages of *Trichomycterus* from the highlands of the Pakaraima Mountains. The upper portions of the study rivers have been connected either through surface flow or by stream capture. Both processes have occurred on multiple time scales and are independent of the patterns seen in the lowlands.

## INTRODUCTION

The Pakaraima Mountains run along the borders of Guyana, Brazil, and Venezuela. These ancient mountains have been the subject of diverse lore of the indigenous inhabitants of the region as well as the western world. Sir Arthur Conan Doyle (1912), in *The Lost World*, imagined dinosaurs and other ancient organisms on the high plateau and Pixar Animation Studios, in the movie *Up* (Docter & Peterson, 2009), imagined a house perched on top of “Paradise Falls” and an undescribed, endemic species of flightless bird. The artists that designed Paradise Falls recognized that many of the rivers of the region fall off of the escarpment in dramatic waterfalls such as Kaieteur, Amaila, and Orinduik Falls.

Although dinosaurs and large, flightless birds do not appear to be denizens of the Pakaraimas, the streams there hold a high degree of endemism (Alofs, Liverpool, Taphorn, & Bernard, 2014; Armbruster & Taphorn, 2011; Hardman, Page, Sabaj, Armbruster, & Knouft, 2002). The Pakaraimas (Figure 1) are drained to the north by the Mazaruni and Cuyuni rivers, to the east by the Potaro River (Essequibo River drainage), to the southwest by the Ireng and Uraricoera rivers (Amazon River drainage), and to the west and northwest by the Caroni River (Orinoco River drainage). The mountains are the remains of Archaean and Proterozoic rocks whose lighter sediments have eroded to fill formerly lacustrine basins such as the Venezuelan Llanos and the Rupununi Savanna of Guyana (see Lujan & Armbruster, 2011, for review). This erosion has left behind a durable core that often has steep faces that the rivers run off of in spectacular waterfalls. Below the falls, the rivers often have some rapids complexes, but quickly reach lowland conditions (Lujan & Armbruster, 2011).

**Figure 1.**
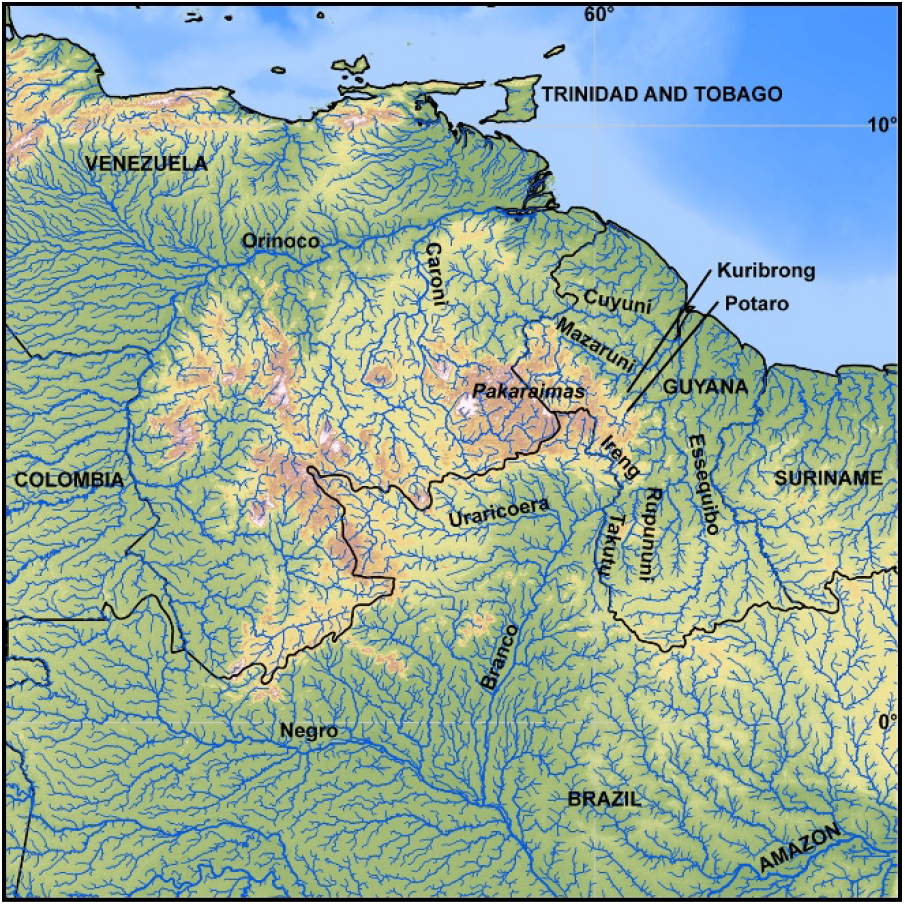
A topographical map of the Pakaraima highlands depicting the physiography of rivers. Major rivers are labelled along their flow. Country names are listed horizontally in all capital letters

Current evidence for the relationships of the rivers draining the Pakaraima Mountains involves, in part, the development and subsequent fragmentation of a paleo-river drainage called the proto-Berbice (Lujan & Armbruster, 2011; Schaefer & do Vale, 1997; Sinha, 1968). The proto-Berbice contained what are now tributaries of the upper rio Branco (Amazon drainage, including the Ireng), the upper Essequibo, the Berbice, and parts of the Courantyne and Orinoco. Meanwhile, the middle and lower Essequibo (including the Potaro/Kuribrong) likely joined the Mazaruni and Cuyuni near where the current mouths are. Slowly, the Amazon River has been capturing streams from the proto-Berbice in an east-west manner. This pattern would suggest a similarity between the faunas of the Potaro/Kuribrong and the upper Mazaruni with the Ireng being more distantly related as it appears to have never been connected into the middle and lower Essequibo + Mazaruni.

However, the upper courses of the rivers have not been explored biogeographically. The likely complex relationships of the upper courses of the rivers were suggested by the description of the crenuchid *Apareiodon agmatos*, and the loricariid taxa *Paulasquama callis, Neblinichthys brevibraccium*, and *N. echinasus* in the upper Mazaruni (Armbruster & Taphorn, 2011; Taphorn, Armbruster, López-Fernández, & Bernard, 2010; Taphorn, López-Fernández, & Bernard, 2008), all of which share affinities with the Orinoco River basin. Given the absence of these taxa in lowland streams, it is likely that these highland taxa were moving via stream capture or other events that connected these highland tributaries. Thus far, the relationships of the highland regions have been scarcely explored systematically. Lujan et al. (2018) found that *Paralithoxus bovallii* (Loricariidae) from the Ireng was more closely related to an undescribed species in the Courantyne than one from the lower Potaro in support of the proto-Berbice hypothesis; however, *Paralithoxus* is not found elsewhere in the Pakaraima highlands. Lujan et al. (in press) found that *Corymbophanes* (Loricariidae), an upper Potaro/Kuribrong endemic, was sister to an undescribed genus from the upper Ireng with the two clades separated by long branch lengths suggesting an ancient relationship.

Coupled with the lack of basic information on the fauna of the region, the area is also under extreme threat by gold and diamond mining with a strong potential of mining eliminating species before they are even discovered (Alofs et al., 2014). In this study, we explore the potential interconnectedness of the high Pakaraima streams by examining the relationships of the pencil catfishes of the genus *Trichomycterus* in order to identify pertinent diversity and to uncover biogeographic patterns that could be duplicated in other Pakaraima organisms.

Trichomycteridae represents a diverse family of freshwater catfishes distributed across the Neotropics. Of the more than 300 recognized species (Fricke, Eschmeyer, & van der Laan, 2018), the majority of species (219) are found in the Trichomycterinae, which contains the genera *Bullockia*, *Cambeva, Eremophilus*, *Hatcheria*, *Ituglanis*, *Rhizosomichthys*, *Scleronema*, *Silvinichthys*, and *Trichomycterus*. Most of the diversity within Trichomycterinae can be attributed to *Trichomycterus*, with all other genera except *Ituglanis* (28 species), *Cambeva* (25 species), *Silvinichthys* (seven species), and *Scleronema* (three species) being monotypic (Fricke et al., 2018). While other genera exhibit apomorphic specializations, the lack of specializations unique to *Trichomycterus* has long made researchers suspect, and later confirm with molecular studies, the non-monophyly of the genus (Baskin, 2016; de Pinna, 2016; Henschel, Mattos, Katz, & Costa, 2018; Katz et al., 2018; Ochoa et al., 2017).

The emerging phylogenetic pattern matches those of other similarly distributed fishes, such as doradid catfishes, characins, and armored catfishes, where distinct clades are geographically linked to a Trans-andean/Amazonian distribution or to south Atlantic coastal drainages (Katz et al., 2018; Ochoa et al., 2017; Ribeiro, 2006). Katz et al. (2018) attempted to solve some of the taxonomic problems of the Trichomycterinae by restricting *Trichomycterus* to a clade that contained the type species (south Atlantic coastal drainages), describing *Cambeva* for a clade sister to *Scleronema*, a clade that is sister to *Trichomycterus sensu stricto*, and referring the Andean, Pagaonian, Amazonian, and Guiana Shield species to “*Trichomycterus*” in quotation marks. “*Trichomycterus*” is paraphyletic and part of a clade that includes *Bullockia*, *Eremophlius sensu stricto*, and *Ituglanis*. Results were similar to those in Ochoa et al. (2017). These patterns are not surprising, given the tectonic and geologic history of the continent that highlights the importance of the Guiana and Brazilian Shields as original uplands of South America, formation of the Andes, and uplift of the Eastern Cordillera (to name a few) with shaping the biogeography of neotropical fishes (Lujan & Armbruster, 2011; Lundberg et al., 1998; Ribeiro, 2006). For ease, we will not be referring to *Trichomycterus* in quotation marks.

*Trichomycterus* are long, slender catfishes generally found only in swift waters. Such habitat, even in the mountains, is patchy, and we suspect that the fishes would be more likely to be isolated to drainages. Recent collections from this region have identified all specimens as *T. guianensis* (Eigenmann, 1912), but we noted significant differences in color and morphology in samples that we have made. Preliminary external visual examinations indicate the possibility for unrecognized diversity and perhaps misidentification of *T. guianensis* in the rivers of this region. The only other species recognized in the region is *T. conradi* (Eigenmann, 1912), and we have found some specimens from the Ireng and Kuribrong rivers that correspond to this species.

Recent studies have illuminated the need to identify unrecognized diversity within *Trichomycterus* and have highlighted the important role that geology and topography play in contributing to that diversity (Katz et al., 2018; Ochoa et al., 2017; Unmack, Bennin, Habit, Victoriano, & Johnson, 2009). In this study, we 1) confirm the discovery of multiple endemic *Trichomycterus* species in the region, 2) examine the diversity and endemism of *Trichomycterus* in the Pakaraima Mountain region with respect to the unique geologic features that have likely influenced their genetic structure, and 3) provide clarification for the identification of *T. guianensis* and *T. conradi* based on morphology.

## METHODS

### Taxon Sampling, DNA Extraction and Sequencing

Collections ranged across multiple years with research permits from Environmental Protection Agency of Guyana as follows, listed as year, reference number: 2011, 030510 BR 130; 2008, 300408 SP: 004; 2014, 040414 SP: 003; 2015, 123115 BR031; 2016, 012016 SP: 003. Fish were either collected with six-foot by ten-foot nylon coated seines with ⅛” mesh, or we joined fishing expeditions of the Patamona who used hiari, a root native to the area around the collection site and a natural source of rotenone (Figure 2). After capture, fish were euthanized in a solution of tricaine methanesulfonate (MS-222) until no sign of respiration was observed for five minutes. Tissue samples were taken from the right pectoral fin or right axial musculature and placed into 1.5 mL vials containing RNALater or ethanol for preservation. Once tissue samples were taken, voucher specimens were fixed in a 3.7% formaldehyde solution for seven days, then rinsed in water for three days, and finally stored in 70% ethanol. Vouchers and tissue samples were deposited in the AUM Fish Collection. Additional materials not collected by the authors were requested from the Royal Ontario Museum (ROM, Table 1).

**Table 1.** Collection information for *Trichomycterus* species used in this study. GenBank Accession numbers are provided for each gene and individual.

| Tissue Catalog | Species ID | Voucher Number | Latitude | Longitude | 16S | COI | Cytb | RAG2 |
| --- | --- | --- | --- | --- | --- | --- | --- | --- |
| AUFT10166 | Ireng, spotted | 67129 | 5.08955 | -59.97514 | Y | Y | Y | Y |
| AUFT10168 | Ireng, spotted | 67129 | 5.08955 | -59.97514 | Y | Y | - | Y |
| AUFT10169 | Ireng, spotted | 67129 | 5.08955 | -59.97514 | Y | Y | Y | Y |
| AUFT10170 | Ireng, spotted | 67129 | 5.08955 | -59.97514 | Y | Y | Y | Y |
| AUFT10212 | T. conradi | 67138 | 5.04398 | -59.97717 | Y | Y | Y | Y |
| AUFT10213 | T. conradi | 67138 | 5.04398 | -59.97717 | Y | Y | Y | - |
| AUFT10234 | Ireng, spotted | 67154 | 5.08388 | -59.98762 | Y | Y | Y | Y |
| AUFT10276 | Ireng, spotted | 67179 | 5.04398 | -59.97717 | Y | Y | - | Y |
| AUFT10294 | T. conradi | 67194 | 5.08867 | -59.96952 | Y | Y | Y | - |
| AUFT10310 | Ireng, spotted | 67172 | 5.08955 | -59.97514 | Y | Y | Y | Y |
| AUFT2110 | T. guianensis | 63677 | 5.30181 | -59.89838 | Y | Y | - | - |
| AUFT2186 | T. cf. guianensis | 62902 | 5.40532 | -59.5439 | Y | Y | - | Y |
| AUFT4743 | T. cf. conradi | 51758 | 4.767118 | -54.56462 | Y | Y | Y | - |
| AUFT6563 | T. guianensis | 62932 | 5.30181 | -59.89838 | - | Y | - | Y |
| AUFT6596 | Potaro, elongate | 62949 | 5.304 | -59.89819 | Y | - | - | Y |
| AUFT6597 | Potaro, elongate | 62949 | 5.304 | -59.89819 | Y | - | - | Y |
| ROMT06183 | Mazaruni, plain | 83791 | 5.4755 | -60.77967 | Y | Y | Y | Y |
| ROMT06184 | Mazaruni, plain | 83791 | 5.4755 | -60.77967 | Y | Y | Y | - |
| ROMT06185 | T. cf. guianensis | 83790 | 5.4755 | -60.77967 | Y | Y | Y | - |
| ROMT06186 | T. cf. guianensis | 83790 | 5.4755 | -60.77967 | Y | Y | Y | - |
| ROMT12696 | T. cf. guianensis | 89932 | 4.95407 | -59.85882 | Y | Y | Y | Y |
| ROMT15527 | T. cf. guianensis | 91392 | 5.272085861 | -59.7026908 | Y | Y | Y | Y |
| ROMT15575 | T. cf. guianensis | 91500 | 5.3759978 | -59.5472803 | Y | Y | Y | Y |
| ROMT15595 | T. conradi | 91436 | 5.413958782 | -59.470252 | Y | Y | Y | Y |

**Figure 2.**
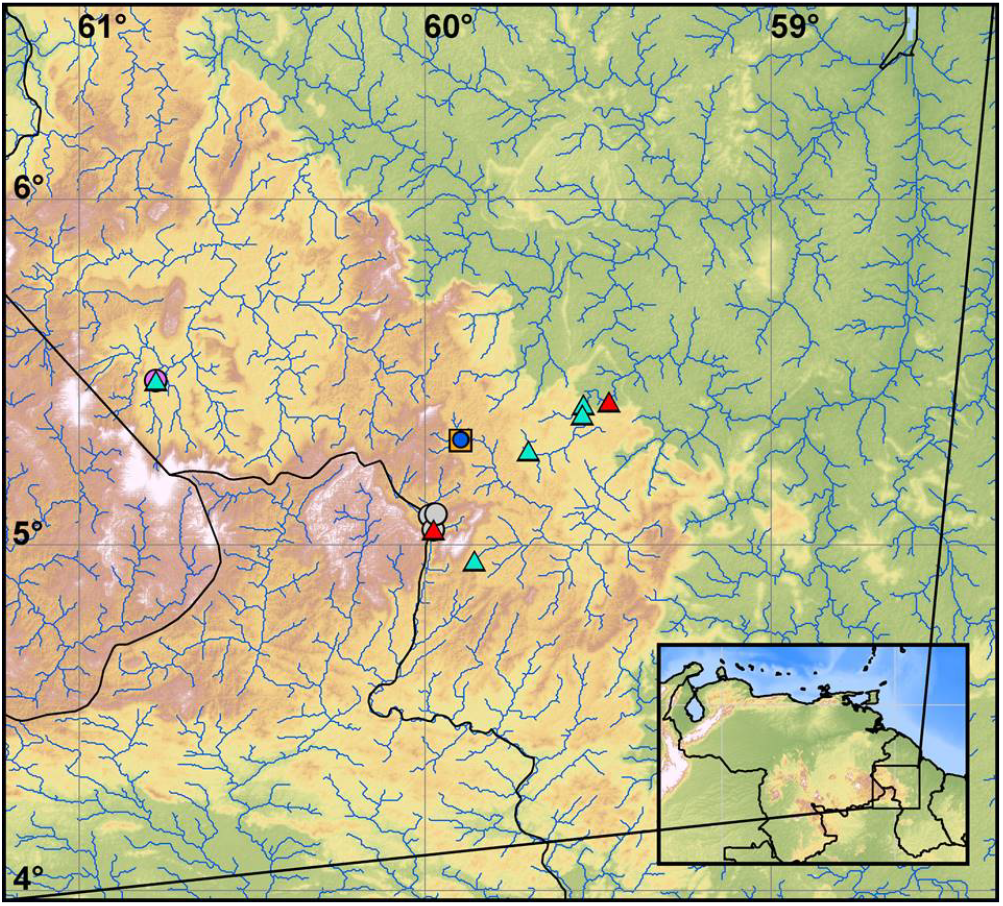
Collection localities for species of *Trichomycterus* found in this study. Color codes correspond to images in Figure 7 and are as follows: red triangles, *T. conradi*; blue circle, *T. guianensis*; teal triangles, *T.* cf. *guianensis*; purple circle, Mazaruni, plain form; orange square, Potaro elongate; gray circle, Ireng, spotted form.

Whole genomic DNA was extracted from tissues using either Chelex or an E.Z.N.A Tissue DNA Kit (Omega BioTek, Norcross, GA). The four genes 16S, COI, cytb, and RAG2 were amplified through 25μL polymerase chain reactions using primers described in Ochoa et al. (2017). The 16S gene was amplified using the following protocol: initial denaturation step of 180 s at 94°C then 30 cycles of denaturation (45 s at 95°C), annealing (30 s at 54°C), and extension (60 s at 68°C) followed by a final extension of 600 s at 68°C. The COI gene was amplified using the following protocol: initial denaturation step of 180 s at 94°C then 30 cycles of denaturation (45 s at 94°C), annealing (30 s at 54°C), and extension (60 s at 68°C) followed by a final extension of 60 s at 68°C. The cytb gene was amplified using the following protocol: initial denaturation step of 180 s at 94°C then 30 cycles of denaturation (45 s at 95°C), annealing (30 s at 54°C), and extension (60 s at 68°C) followed by a final extension of 60 s at 68°C. The RAG2 gene was amplified using a two-step protocol. The first reaction was performed using the touchdown protocol described in Lovejoy & Collette (2001) with RAG164F and RAG2R6 primers. The second PCR used 1.5μL of template from the first run and primers 176R and RAG2Ri under the following conditions: initial denaturation step of 30 s at 95°C then 35 cycles of denaturation (30 s at 95°C), annealing (45 s at 56°C), and extension (90 s at 72°C). Primers used for PCR amplification were also used for DNA sequencing for all genes, with 176R and RAG2Ri used for sequencing RAG2.

The products were visualized and size-verified on a 0.8% agarose gel. PCR purification, sample preparation, and Sanger sequencing were performed at GeneWiz (South Plainfield, NJ). Chromatographs from forward and reverse reads were imported into Geneious v. 10.2.3 (Kearse et al., 2012) for assembly. Assembled contiguous sequences were aligned using the MUSCLE algorithm (Edgar, 2004), and results were checked by eye. Due to length variation among sequences generated in this study and those of Ochoa et al (2017), alignments were trimmed to the following lengths: 16S: 466; COI 522; cytb: 858; myh6: 543; and RAG2: 885. Each individual gene tree was analyzed with *Scleronema minutum* as an outgroup, while the concatenated dataset (3579bp) included members from Ochoa et al’s (2017) clades D1, D2, D3, and E with *S. minutum* as an outgroup. Data were exported both as individual alignments and as a concatenated dataset for phylogenetic analysis.

### Phylogenetic Analysis

Best-fit models of evolution were tested using PARTITIONFINDER2 (Lanfear, Frandsen, Wright, Senfeld, & Calcott, 2017). Models were tested on individual gene trees, and then on the concatenated dataset. The resulting data blocks were then used in Bayesian Inference analysis. Bayesian Inference was performed using MrBayes v. 3.2.6 on XSEDE via CIPRES Science Gateway (Miller, Pfeiffer, & Schwartz, 2010). Each dataset had 2 runs with 4 chains run for 15 million generations, sampling once every 1,000 generations. The parameters and trees were summed in MrBayes v. 3.2.6 using the default 25% burn-in. The resulting 50% majority consensus rule phylogeny is reported.

### Maps

The maps produced for this paper were created in ARCGIS, ARCMAP V. 10.3.1; ESRI, 2011). Digital elevation models and rivers are from HydroSHEDS by the United States Geological Service and World Wildlife Federation (https://www.worldwildlife.org/pages/hydrosheds; https://hydrosheds.cr.usgs.gov/). Width of rivers is by Strahler number (stream order).

Bathymetry is ETOPO1 from the National Geophysical Data Center (https://www.ngdc.noaa.gov/mgg/global/). Color ramps for elevation and bathymetry are from Environmental Systems Research Institute’s Color Ramps v. 3.0 (https://www.esri.com/arcgis-blog/products/product/imagery/esri-color-ramps-version-3-0/). Country borders and graticules are from Natural Earth (https://www.naturalearthdata.com/).

## RESULTS

Four genes trees were analyzed separately, then combined into a concatenated analysis. The first individual gene tree is cytochrome b (cytb, Figure 3), which results in a well-supported clade of Pakaraima *Trichomycterus*. Members of true *Trichomycterus guianensis* are found sister to the Potaro, elongate form. This clade is sister to another well-supported clade of *T. cf. guianensis* + Mazaruni, plain form. These are all sister to a single representative of the *T. conradi*. This analysis did not include the Ireng, spotted form that is present in other analyses.

**Figure 3.**
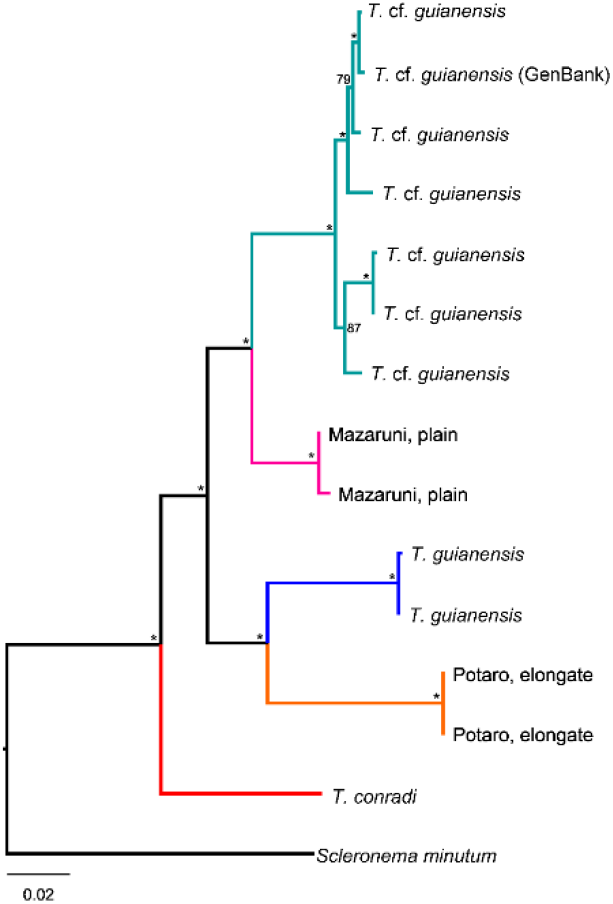
Fifty per cent majority rule consensus tree from Bayesian inference of cytochrome b sequences. Nodes labelled with an asterisk (*) indicate posterior probabilities >90%. Values less than 90% are written on the trees. Branches are colored to match localities as seen in Figure 1. Tip labels correspond to individuals as denoted in Table 1.

The second gene tree generated from our data is based on COI (Figure 4). This analysis again places *T. cf. guianensis* sister to the Mazaruni, plain form. In contrast to the cytb phylogeny, this clade is sister to the *T. conradi*; however, this relationship is weakly supported. The *(T. cf. guianensis* + Mazaruni, plain) *T. conradi* clade is sister to another clade consisting of the Ireng, spotted form and true *T. guianensis.* The interrelationships among the clades are poorly supported, but each recognized morphotype is well-supported with the exception of the Ireng, spotted form. Finally, the Potaro, elongate form is missing from this analysis. Overall, the COI tree is much less resolved than the other trees, with some nodes not reaching 90% posterior probability.

**Figure 4.**
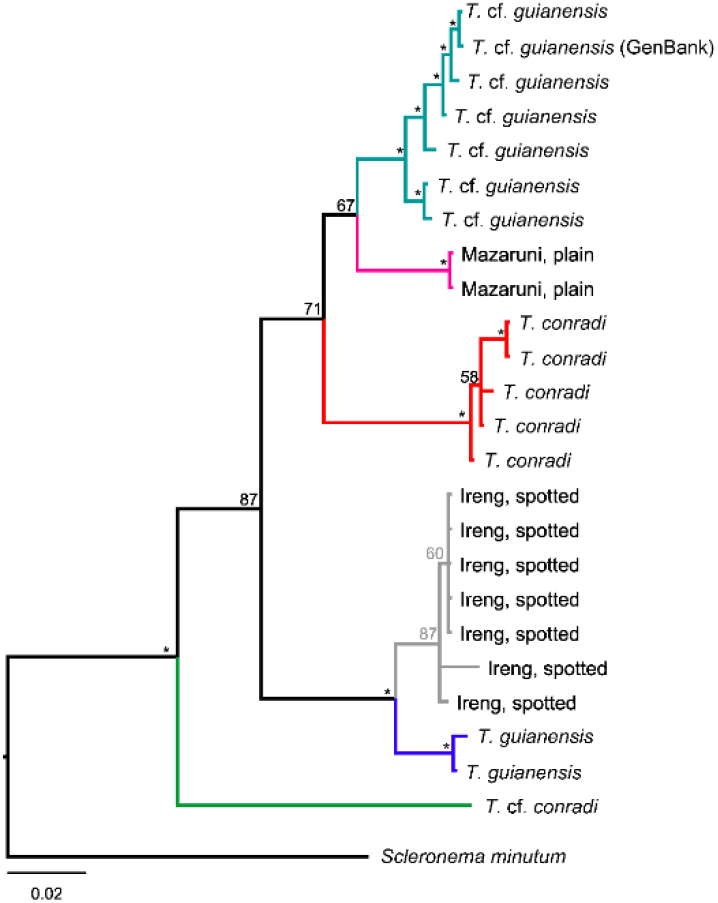
Fifty per cent majority rule consensus tree from Bayesian inference of COI sequences. Nodes labelled with an asterisk (*) indicate posterior probabilities >90%. Values less than 90% are written on the trees. Branches are colored to match localities as seen in Figure 1. Tip labels correspond to individuals as denoted in Table 1.

Ribosomal 16s data place *T. cf. guianensis* sister to the Mazaruni, plain form (Figure 5). This clade is sister the *T. conradi*. As seen in the COI analysis, despite geographic proximity, the Ireng, spotted form is sister to true *T. guianensis* rather than the *T. conradi*. The Potaro, elongate form is sister to the rest of the member clade.

**Figure 5.**
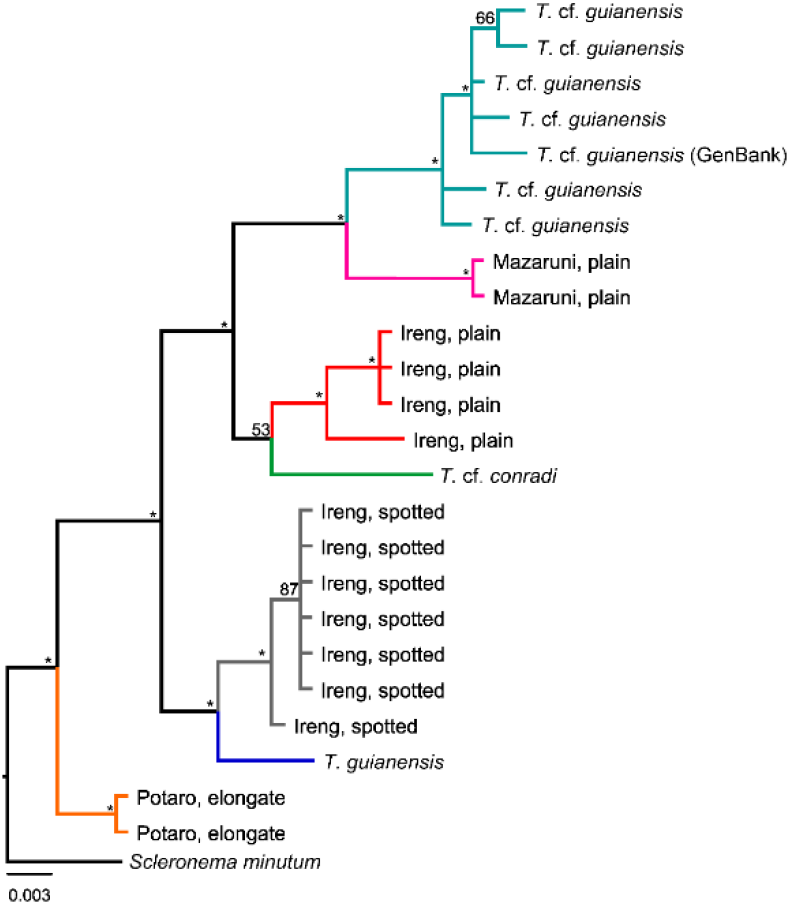
Fifty per cent majority rule consensus tree from Bayesian inference of 16S sequences. Nodes labelled with an asterisk (*) indicate posterior probabilities >90%. Values less than 90% are written on the trees. Branches are colored to match localities as seen in Figure 1. Tip labels correspond to individuals as denoted in Table 1.

Nuclear DNA analysis from the RAG2 data was the most divergent from the remainder of the data (Figure 6). Again, *T. cf. guianensis* is recovered sister to the Mazaruni, plain form, but this is the only similarity with the other gene trees. The RAG2 data show true *T. guianensis* sister to the Ireng, spotted form. They form a clade sister to the Potaro, elongate form. The *T. conradi* is paraphyletic and its relationships are unresolved due to a polytomy.

**Figure 6.**
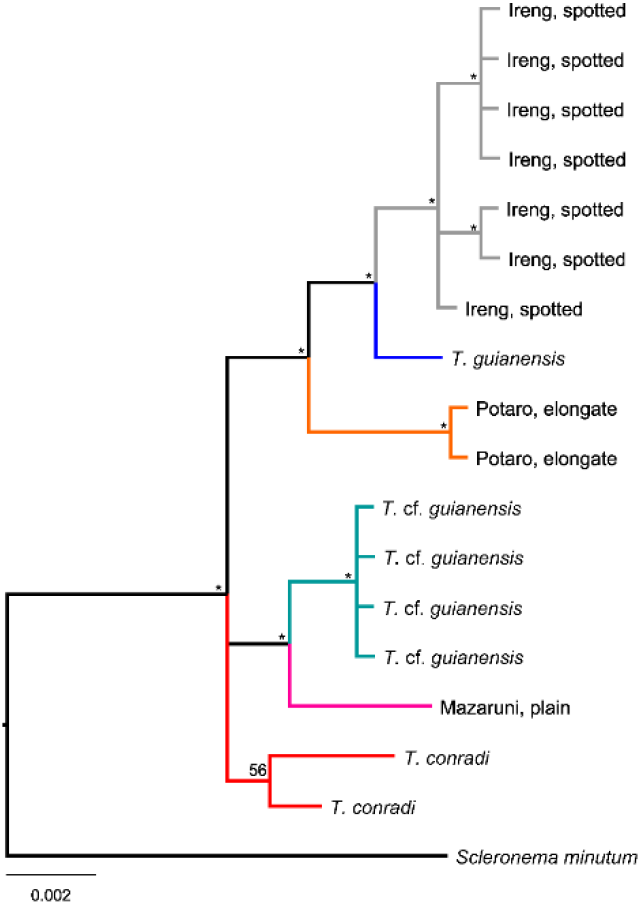
Fifty per cent majority rule consensus tree from Bayesian inference of RAG2 sequences. Nodes labelled with an asterisk (*) indicate posterior probabilities >90%. Values less than 90% are written on the trees. Branches are colored to match localities as seen in Figure 1. Tip labels correspond to individuals as denoted in Table 1.

With gene tree heterogeneity rampant in this analysis, the four genes were concatenated and analyzed with the D1, D2, D3, and E clades from Ochoa et al (2017, Figure 7). This tree, with 24 individuals from our analysis, shows that all morphotypes we identified *a priori* are monophyletic. Two distinct clades compose Pakaraima *Trichomycterus*: *Trichomycterus cf. guianensis* + Mazaruni, plain form are sister to the *T. conradi*. This clade is sister to another clade consisting of *Trichomycterus guianensis +* Ireng, spotted which are sister to the Potaro, elongate form. Each of these relationships are supported with >90% posterior probability, while deeper relationships remain unresolved.

**Figure 7.**
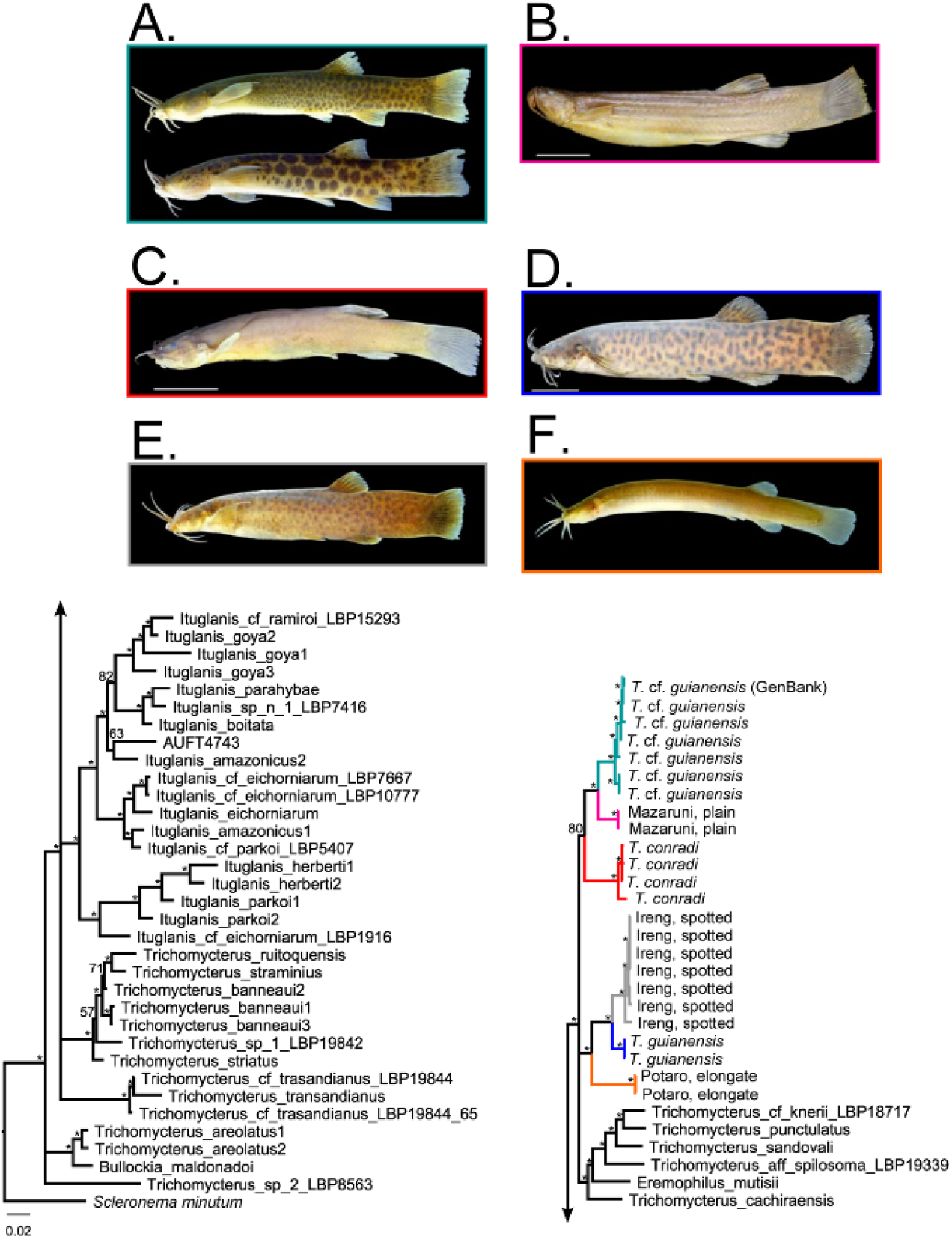
Fifty per cent majority rule consensus tree from Bayesian inference of concatenated sequences. Nodes labelled with an asterisk (*) indicate posterior probabilities >90%. Values less than 90% are written on the trees. Branches are colored to match localities as seen in Figure 1. Tip labels correspond to individuals as denoted in Table 1. These sequences are combined with the D1, D2, D3, and E clades from Ochoa et al (2017). Arrows connect disconnected branches in the phylogeny. Outlines of the photographs of specimens correspond to clade color and symbol color in Figure 1. A. *T.* cf. *guianensis*. B. Mazaruni, plain form. C. *T. conradi*. D. true *T. guianensis*. E. Ireng, spotted form. F. Potaro, elongate form.

## DISCUSSION

Our results demonstrate the presence of multiple species of *Trichomycterus* in the Pakaraima Mountains of Guyana. There were two major clades, one consisting of *T. guianensis* and two undescribed species, and the other of *T. conradi* and two undescribed species. Both *T*. *guianensis* and *T. conradi* appear to be the rarer species in the region (based on collections), and *T*. *guianensis* is not as widespread as previously believed. Based on examination of the types and comparison with specimens we have collected, *Trichomycterus guianensis* is a deep bodied species with irregular blotches (Figure 7D). *Trichomycterus* sp. Ireng Spotted (Figure 7E) is the dominant species in the Ireng, and it is similar in morphology to *T. guianensis* and was recovered as sister to it. In morphology, the Ireng Spotted species is even shorter and deeper-bodied than *T. guianensis*. Sister to the clade of *T. guianensis* and the Ireng Spotted species is a very elongate, almost entirely brown species from near Ayangana in the far upper Potaro with miniscule pelvic fins (*Trichomycterus* sp. Potaro Elongate, Figure 7F). It was found in sluggish, swampy areas, which is habitat more indicative of *Ituglanis*, but the species lacks the diagnostic characters of *Ituglanis* (JWA pers. obs.), and we found it to be a member of *Trichomycterus* in the molecular analysis. *Trichomycterus* sp. Potaro Elongate and *Trichomycterus* sp. Ireng Spotted are at the opposite extreme of *Trichomycterus* morphology, suggesting a strong capacity for body rearrangement in the genus.

The other major clade contains a wide-spread, elongate species with dark, small or large, regular spots (*Trichomycterus* cf. *guianensis*, Figure 7A). This species is found in the upper Potaro, Kuribrong, and Mazaruni rivers. Little geographic structure was present in the specimens examined, suggesting fairly recent movement between the basins. Sister to this species is a similar but unspotted species from the Mazaruni River (*Trichomycterus* sp. Mazaruni Plain, Figure 7B). Finally, sister to the other two species is a clade that consists of a few individuals from the Ireng as well as a specimen from the lower Kuribrong of a plain-colored species that appears to be *T. conradi* (Figure 7C).

Two species of *Trichomycterus* have been described from the upper Caroni, Orinoco River drainage section of the Pakaraima Mountains: *T. celsae* Lasso and Provenzano 2002 and *T. lewi* Lasso and Provenzano 2002. The Orinoco species do appear to be different from the species analyzed here, and it appears that there are additional undescribed species from that region. Unfortunately, we do not have tissue samples from these species.

### Biogeography of the Pakaraima Mountains

The biogeographic story that the species of *Trichomycterus* of the Pakaraimas tell is a complex one. *Trichomycterus* cf. *guianensis* appears to have moved between river systems relatively easily. Mazaruni samples are sister to those in the Kuribrong and Potaro rivers, but the Mazaruni samples are paraphyletic. Tributaries of the Mazaruni interdigitate with the Kuribrong and Potaro rivers, and species living as high in their drainages as *Trichomycterus* would be more likely to be able to move via river capture events where tributaries erode their divides and switch from one system to the next. Anecdotal reports suggest that the upper courses of at least the Potaro and Kuribrong connect during particularly rainy times; flying over the area reveals numerous fissures that seem to run between the two rivers (JWA pers. obs.). These drainages also interdigitate with Caroni and Ireng tributaries. Some of the specimens from the Caroni do appear similar to the elongate, spotted species of Guyana, but we did not find anything similar in the Ireng despite extensive searching.

The upper Caroni and the Ireng were once part of the proto-Berbice paleodrainage basin along with the upper Branco, upper Essequibo, Berbice, and Courantyne rivers while the Mazaruni was likely independent (Lujan and Armbruster, 2011). The Essequibo makes a westward bend near Massara and away from a nearby Berbice tributary (Gibbs & Barron, 1993), suggesting a likely point of demarcation between the upper Essequibo as part of the proto-Berbice and the lower Essequibo, which probably joined with the Mazaruni at the present mouth of the Essequibo. This would mean that the Potaro and Mazaruni were part of the same system and not part of the proto-Berbice. However, the mixing of Ireng, Potaro, and Mazaruni *Trichomycterus* in the phylogeny suggests that there likely existed faunal exchange between the proto-Berbice, Potaro, and Mazaruni rivers at least in the highlands prior to the breakup of the proto-Berbice during the Pliocene and Pleistocene potentially leading to complex interrelationships between these basins. A similar finding was made in Lujan et al. (in press) who found that *Corymbophanes* was sister to a new genus from the Ireng; however, the branch lengths were much longer than what was observed here. Further exploration into the relationships of *Trichomycterus* along with a molecular clock will likely lead to fascinating insights into the biogeography of the Pakaraima Mountains, but this further insight will require extensive collecting in the difficult to explore Brazilian tributaries of the Pakaraima Mountains and further collecting in Venezuela, that is difficult now because of civil strife.

*Trichomycterus conradi* appears to be a more lowland form found in the rapids below Kaieteur and Amaila Falls on the Potaro and Kuribrong, respectively, as well as in the Ireng. The shallow nodes between the Kuribrong and Ireng samples sequenced suggest that movement has been relatively recent. We were only able to obtain 16S sequences for a specimen of *T. cf. conradi* from the Maroni River of eastern Suriname, and it was sister to *T. conradi*. A similar distribution across the northern Guiana Shield was found for *Paralithoxus bovallii* from the Ireng River and hypothesized new species related to it in the Potaro, Courantyne, and Coppename rivers (Lujan et al. 2018). The distributions of *T. conradi* and *P. bovallii sensu lato* suggest interconnectivity across the Guiana Shield even for small fishes restricted to fast-flowing streams. Clearly, we are just beginning to understand the complexities of the biogeography of the western Guiana Shield and the interconnectedness of it with the eastern portion of the shield.

### Threats to Biodiversity in the Guyana Highlands

The Pakaraimas represent the cores of ancient mountains, which are among the main sources of gold and diamonds. Alofs et al. (2014) review some of the issues with gold mining in the upper Mazaruni River, and we have observed similar issues in the Kuribrong and Potaro Rivers as well. Large swaths of forest have been removed from around the rivers with the sediment pumped through sieves to extract gold and diamonds. Gold is removed with mercury amalgamation leading to high mercury levels in the water, fishes, and humans (Miller et al. 2003) and large swaths of forest replaced by denuded landscapes and toxic spoil ponds. On larger rivers like the lower Potaro, large dredging machines suck up sediment and process it directly in the river leaving behind piles of gravel in the river that alter the natural hydrology. Although Hardman et al. (2002) did not find significant differences between their study of the fishes of the Potaro River and Eigenmann (1912), certain species that had been present and common in Eigenmann’s survey were absent 90 years later. Mol & Ouboter (2004) and Brosse, Grenouillet, Gevrey, Khazraie, & Tudesque (2011) found that the erosion related to gold mining has reduced fish diversity. As of our 2014 trip to the upper Kuribrong and 2016 trip to the Ireng, there was little impact to the rivers from mining; however, a recently completed road now provides easier access to the upper Kuribrong, and one small mine was observed. The lower Kuribrong has been heavily impacted, and after flying over the Potaro River in 2014, JWA can state that the Potaro looks less clear than it had during the 1998 expedition reported in Hardman et al. (2002).

As expressed by Alofs et al. (2014) for the upper Mazaruni, the whole high plateau of the Pakaraimas supports an endemic fauna as is evidenced here. Although there is some interconnectivity of the river systems, narrow endemic *Trichomycterus* are found in each of the rivers in this study. Conservation of this unique landscape that has become part of our shared cultural heritage is important, and further studies on the unique fauna of the region are needed.

## ACKNOWLEDGEMENTS

This research was supported by an undergraduate Research Grant-In-Aid from the Department of Biological Sciences, Auburn University to HJP and a grant from the COYPU Foundation to JWA. JWA and DCW would like to thank Donald Taphorn, Nathan Lujan, and Elford Liverpool as well as Aiesha Williams and Chuck Hutchinson of the Guyana WWF for arranging trips to collect fishes in Guyana, and Ovid Williams for acting as a liaison to indigenous communities. We would like to thank the Patamona people of Kaibarupai, Ayangana, and Chenapowu for hosting us on expeditions and imparting their knowledge and skills in the field. Numerous people aided in the collection of specimens, and we owe them our deepest gratitude. Special thanks to Liz Ochoa for helping us design and implement this study. Thanks to Erling Holm for sending specimens and tissues from ROM.

## DATA ACCESSIBILITY STATEMENT

Sequence data are available on GenBank. Accession numbers for each specimen and gene are listed in Table 1.

*Note: Accession numbers will be included upon acceptance of the paper*.

